# Single-gene resolution of diversity-driven community overyielding

**DOI:** 10.1101/2022.10.14.512290

**Authors:** Samuel E. Wuest, Lukas Schulz, Surbhi Rana, Julia Frommelt, Merten Ehmig, Nuno D. Pires, Ueli Grossniklaus, Christian S. Hardtke, Ulrich Hammes, Bernhard Schmid, Pascal A. Niklaus

**Affiliations:** Department of Evolutionary Biology and Environmental Studies and Zurich-Basel Plant Science Center, University of Zurich, Winterthurerstrasse 190, 8057 Zurich; Department of Plant and Microbial Biology and Zurich-Basel Plant Science Center, University of Zurich, Zollikerstrasse 107, 8008 Zurich; Department of Geography, Remote Sensing Laboratories, University of Zurich, 8057 Zurich, Switzerland; Agroscope, Group Breeding Research, Mueller-Thurgau-Strasse 8, 8820 Waedenswil, Switzerland; Plant Systems Biology, School of Life Sciences, Technical University of Munich, 85354 Freising, Germany; Department of Plant Molecular Biology, University of Lausanne, Biophore Building, Lausanne 1015, Switzerland; Department of Systematic and Evolutionary Botany, University of Zurich, Zollikerstrasse 107, 8008 Zürich

## Abstract

In plant communities, diversity often increases community productivity and functioning, but the specific underlying drivers are difficult to identify. Most ecological theories attribute the positive diversity effects to complementary niches occupied by different species or genotypes. However, the type of niche complementarity often remains unclear, including how complementarity is expressed in terms of trait differences between plants. Here, we use a gene-centred approach to identify differences associated with positive diversity effects in mixtures of natural *Arabidopsis thaliana* genotypes. Using two orthogonal genetic mapping approaches, we found that between-plant allelic differences at the *AtSUC8* locus contribute strongly to mixture overyielding. The corresponding gene encodes a proton-sucrose symporter and is expressed in root tissues. Genetic variation in *AtSUC8* affected the biochemical activities of protein variants and resulted in different sensitivities of root growth to changes in substrate pH. We thus speculate that - in the particular case studied here - evolutionary divergence along an edaphic gradient resulted in the niche complementarity between genotypes that now drives overyielding in mixtures. Identifying such genes important for ecosystem functioning may ultimately allow the linking of ecological processes to evolutionary drivers, help to identify the traits underlying positive diversity effects, and facilitate the development of high-performing crop variety mixtures in agriculture.

## Introduction

Functional differences between plants are major determinants of the composition, diversity, and functioning of communities (Loreau, 2000; Lavorel and Garnier, 2002; McGill et al., 2006; Plas et al., 2020). Some of these differences represent adaptations of species to sets of environmental conditions, also termed niches (Violle and Jiang, 2009; Roscher et al., 2015). Many theories support the notion that niche complementarity among plants underlies commonly observed positive biodiversity– ecosystem functioning relationships ((Tilman et al., 1996; Hector et al., 1999; Tilman et al., 2006; Reich et al., 2012; Zuppinger-Dingley et al., 2014; Turnbull et al., 2016). While plausible, it currently is less clear how the relevant niche dimensions underlying such functional complementarity can be identified, and how complementarity manifests itself in specific trait differences between plants (Kraft et al., 2015; Crutsinger, 2016; Barry et al., 2019; Plas et al., 2020). An important reason for this knowledge gap is that, rather than quantifying niche space directly, niche complementarity is mostly indirectly implied from observed higher-level phenomena, such as increasing productivity with increasing biodiversity, with little reference to the underlying physiology (Barry et al., 2019; Plas et al., 2020). Furthermore, approaches focusing on traits as surrogates for niches (Roscher et al., 2015) struggle with the problem of co-varying explanatory variables and the difficulty to separate correlation from causation: traits often co-vary because of fundamental evolutionary trade-offs between ecological strategies (Wright et al., 2004; Díaz et al., 2015).

Finally, it also is likely that not a single but many small phenotypic trait differences together determine niche complementarity between plants (Kraft et al., 2015; Montazeaud et al., 2020). The multivariate nature of phenotypic differences associated with niche complementarity thus makes it difficult to pinpoint specific mechanisms that underly biodiversity–productivity relationships (Cadotte, 2017; Huang et al., 2018). Therefore, the question arises whether niche complementarity as manifested in functional trait differences (Roscher et al., 2015) is a phenomenon too complex to be studied using reductionistic experimental methods.

Positive biodiversity–productivity relationships occur not only at the inter-but also at the intra-specific level; for example, mixtures of genotypes of natural plants and crops often overyield relative to monocultures of the same genotypes (see, e.g., Hughes and Stachowicz, 2004; Crutsinger et al., 2006; Kiær et al., 2009; Crawford and Whitney, 2010; Reiss and Drinkwater, 2018), although there are exceptions (Bongers et al., 2020). It is reasonable to assume that the mechanisms underlying niche complementarity and overyielding are similar in both cases, although there is clearly a larger potential for niche differences among species than among genotypes of the same species.

Here, we focus on the study of complementarity among genotypes of the model plant species *Arabidopsis thaliana*. A major advantage of this approach is that the diversity of traits and alleles cannot only be manipulated by assembling communities from an existing pool of genotypes but also through crosses (**Figure 1**). Crosses allow, within the limits of linkage disequilibrium, a redistribution of genetic variation, and therefore trait variation, between genotypes. The assembly of new communities that differ in their genetic composition then allows us to establish causal links between genetic diversity and community-level properties (Wuest and Niklaus, 2018; McGale et al., 2020) (**Figure 1**). Several recently published papers have expanded the traditional approach that links genetic differences amongst individuals to their phenotypic variation to the genetic study of the properties of ecological communities (Wuest and Niklaus, 2018; Wuest et al., 2019; McGale et al., 2020; Turner et al., 2020; Montazeaud et al., 2022). For example, and in analogy to keystone species that exhibit disproportionately large effects on ecosystems, Barbour and colleagues describe a plant “keystone gene” whose presence determined the stability of an experimental trophic food web containing plants, aphids and their parasitoids (Barbour et al., 2022). Together, these publications demonstrate that genetic effects can cascade across layers of increasing biological complexity, sometimes in unexpected ways. Here, we employed a genetic approach to study how genetic *diversity* affects plant community overyielding and combined it with ecological and physiological experiments to investigate the specific type of complementarity.

**Figure 1:**
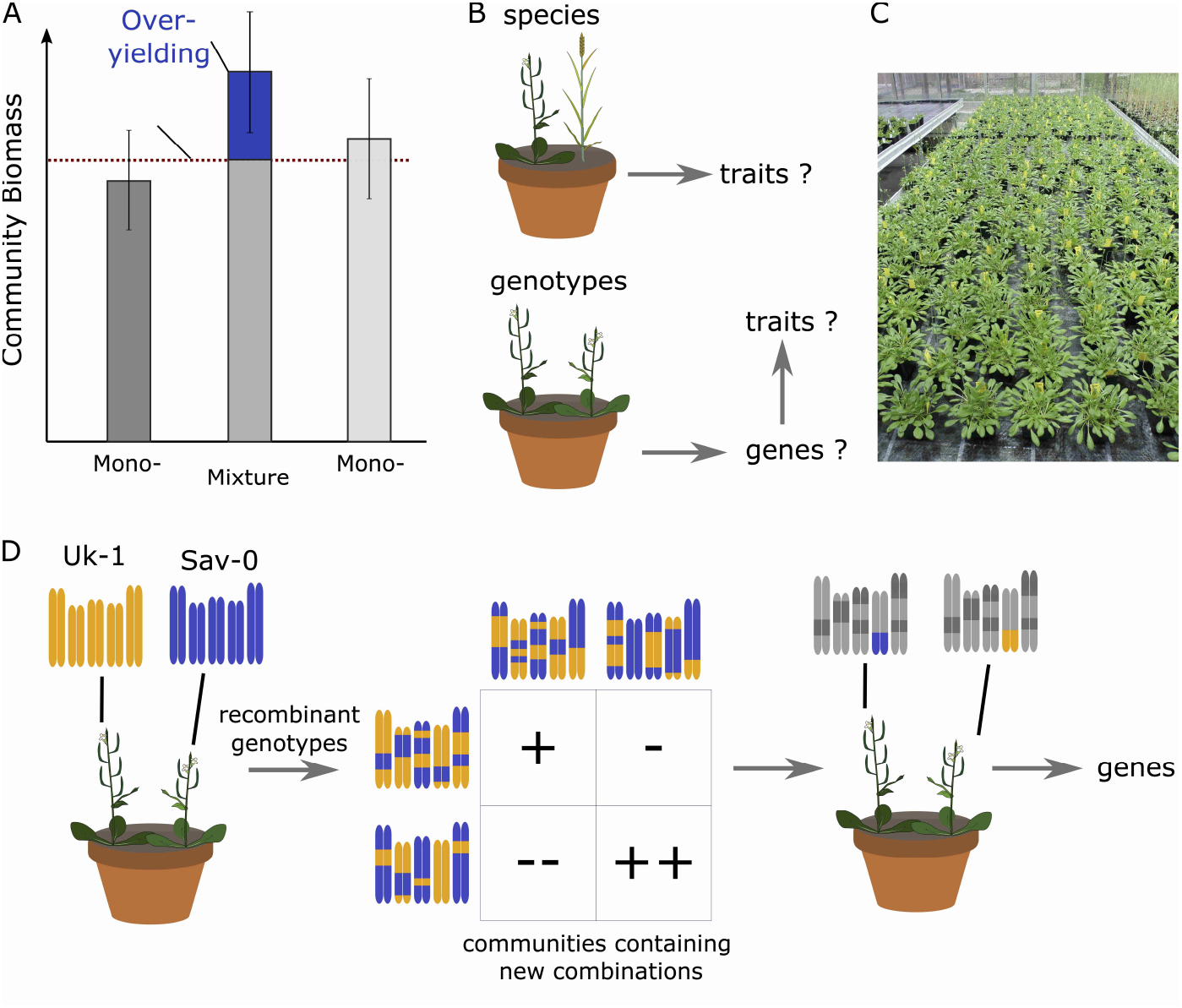
Experimental approaches to the genetic dissection of positive diversity effects. **A**. A positive diversity effect (blue) in pair-wise mixtures denotes the estimated deviation of mixture yield from expectations based on monoculture yields. Estimating this deviation is difficult, because it combines three error terms (two monoculture productivity estimates and one mixture productivity estimate). **B**. Positive effects on productivity can be found with increasing species or increasing genotype diversity within a community. Past work has put much effort into studying the underlying functional trait differences, but our work is concerned with firstly studying the underlying genetic differences, and then trying to infer functional trait differences from genes **C**. Experimental setup used in this study, showing model communities consisting of four plants and different pairwise genotype combinations. **D**. Schematic representation of how a genotypic diversity effects (left; Umkirch-1 + Slavice-0) can be further dissected into genetic diversity effect, by the use of crosses and genetic recombination followed by the assembly of new genotype pairs into model communities. “+” (or “-”) denote community performances that are either higher (or lower) than expected.

## Results

In order to genetically dissect the mechanisms that underly biodiversity effects on productivity, we first needed to identify genotypes that overyield when grown together in mixture, i.e., communities that produce more biomass than the average of their monocultures (**Figure 1 A**). We tested overyielding in communities containing one of ten pairs of *Arabidopsis thaliana* (L.) Heynh. genotypes. (**Supplementary Figure 1 A**). We used these pairs because they are the parents of publicly available recombinant inbred lines, a formidable resource for genetic studies and mapping. Overyielding estimates in this experiment were all not significantly different from zero. This was not unexpected, because overyielding is calculated as difference between three yield values (of the mixture, and the two monocultures); a high replication of all three communities is therefore required to compensate for the error propagation in this calculation. However, model plant communities that contained the two accessions Slavice-0 (Sav-0) and Umkirch-1 (Uk-1) overyielded consistently across three substrates and across different pot sizes. We replicated this effect in a second experiment with two different pot sizes and two plant densities (**Supplementary Figure 1 B**). Across all experimental settings, mixtures of Sav-0 and Uk-1 yielded an average 5.6% more biomass (range: 0–12%) than expected based on monoculture productivities. This effect is relatively large for a pot-based within-species experiment. For comparison, the average overyielding in field trials with crop variety mixtures typically ranges from 2 to 4% (Kiær et al., 2009; Borg et al., 2018; Reiss and Drinkwater, 2018; Kristoffersen et al., 2020).

To overcome the challenges in determining overyielding due to low power resulting from error propagation (**Figure 1 A**), we adopted competition diallels (**Figure 2 A**) (Griffing, 1956; Harper, 1977; Griffing, 1989; Bossdorf et al., 2004). In these, general and specific combining abilities (GCAs and SCAs, **Figure 2 A**) can be taken as proxies for additive and non-additive mixing properties of genotypes and genotype combinations. Here, we used a half-diallel containing 18 randomly selected recombinant inbred lines (RIL) derived from a cross between Sav-0 and Uk-1, and the two parental lines. These RILs had been created to allow the map-based cloning of the BREVIS RADIX (BRX) gene, at which natural variation causes strong root architectural differences between Sav-0 and Uk-1 (Mouchel et al., 2004) - differences that may be expected to drive complementarity in genotype mixtures. The 20 chosen genotypes were now grown in all pair-wise combinations. The diallel was replicated four times, at different dates (temporal blocks). We further used two different substrates (sand-rich and peat-rich soils, two blocks each). We determined the average SCA across the four blocks for each of the 210 community compositions (190 genotype mixtures plus 20 monocultures). To adjust for differences in community productivity between substrates, and to obtain a normal distribution of residuals, we scaled the estimated SCAs by division by the average community biomass on the respective substrate. SCA thus was expressed as effect relative to the mean productivity of all communities on the substrate. Next, we tested if variation in SCA among the different communities could be attributed to genetic differences at specific genomic regions. Since the published marker density for the RIL population used here was relatively low, we first constructed high-resolution genotype maps by whole-genome re-sequencing of each line (Methods, **Supplementary Figure S 2 A**). We then used marker-regression to compare SCAs of communities that were either mono-allelic or bi-allelic at a given marker region, i.e., we tested for effects of allelic diversity. We found that specific combining ability was positively associated with genetic differences at a single quantitative trait locus (QTL) on chromosome 2. The high-density marker map allowed us to resolve this QTL to a very small genomic region, spanning approximately 178 kb (**Figure 2 B**). Mixtures that exhibited allelic diversity in this region exhibited a 2.8% (+/-0.8% s.e.m.) higher SCA than mixtures that contained only one of the two alleles (“mono-allelic” communities, **Figure 2 C**). At the same time, mono-allelic genotype mixtures (mixtures containing only the Sav-0 or only the Uk-1 allele at the identified QTL on chromosome 2, but any allele combination at other loci) had a 0.8% higher SCA than genotype monocultures (no allelic differences at any locus). Therefore, a single QTL on chromosome 2 seems to explain a high proportion of overyielding in Sav-0–Uk-1 genotype mixtures.

**Figure 2:**
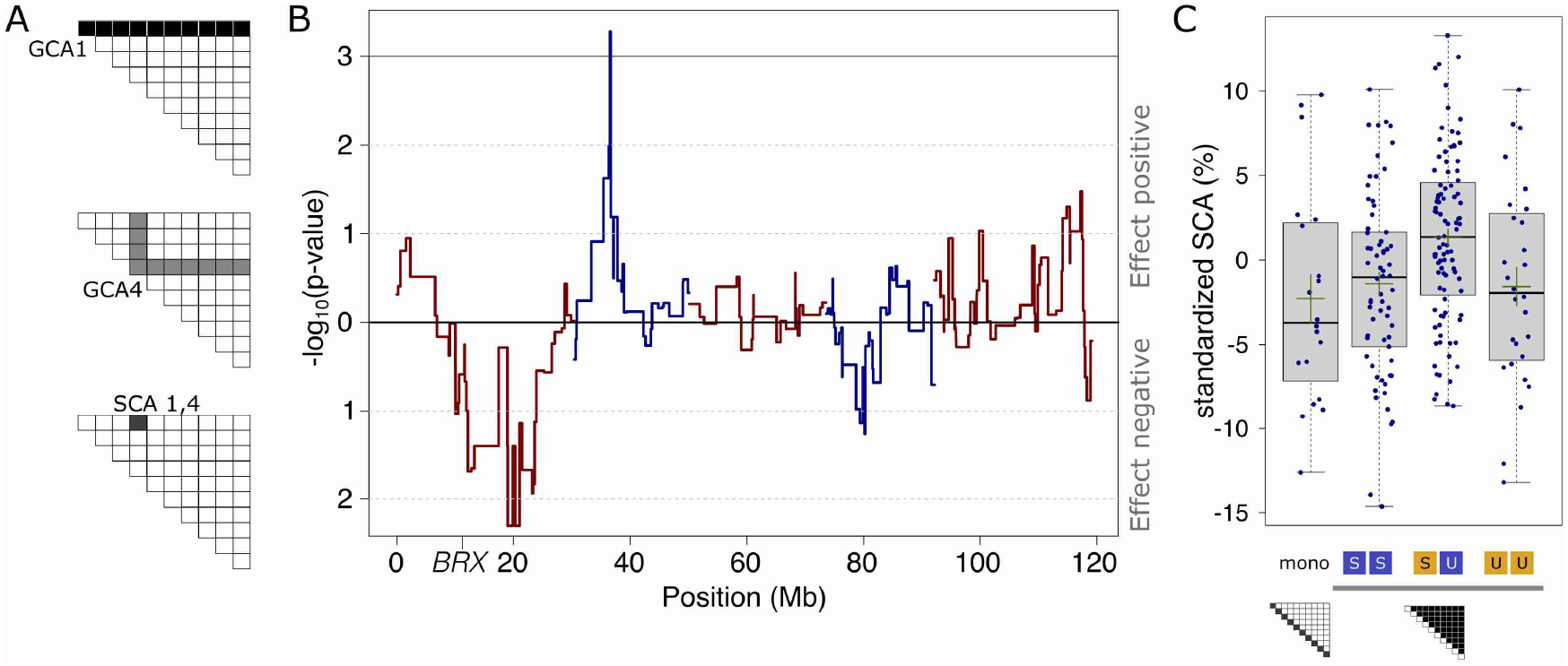
Genotypic and allelic diversity effects. **A**. Illustration of the concepts of General and Specific Combining Abilities (GCA and SCA) derived from genotypic communities assembled according to a competition half-diallel design. GCAs of genotypes 1 and 4 are estimated from productivities of all mixtures in which these genotypes occur, SCA_1,4_ denotes the estimated productivity deviation of communities containing these two genotypes after accounting for GCAs. **B**. QTL map of allelic diversity associated with variation in SCA within genotypic mixtures. Blue and red lines denote the different chromosomes. “BRX” indicates the location of the BREVIS RADIX gene. **C**. Boxplots showing SCA distributions of different communities: genotypic monocultures (mono), genotypic mixtures but allelic monocultures at the QTL on chromosome 2 (SS and UU), genotypic mixtures and allelic mixtures at the QTL (SU). Green lines denote mean values +/-s.e.m. Genotypic mixtures overall exhibit slightly but significantly higher standardized SCAs values than genotypic monocultures (∼ 0 vs −2.7%).

The Uk-1 accession was originally collected from the banks of the Dreisam river in the Schwarzwald of southern Germany. This region is characterized by an edaphic gradient with pH ranging from neutral to strongly acidic (**Supplementary Figure 3**). Previous work has shown that the Uk-1 loss-of-function allele of the *BREVIS RADIX* (*BRX*) gene confers a fitness advantage to plants grown on acidic soil (Gujas et al., 2012) and alters root architecture and plant competition (Mouchel et al., 2004; Shindo et al., 2008). In our experiment, allelic diversity at the *BRX* locus was not associated with community overyielding (**Figure 2 B**, on lower arm of chromosome 1). Nevertheless, we speculated that the observed overyielding might have been driven by niche complementarity that resulted from adaptive divergence along this edaphic gradient. The identified QTL contained 16 protein-coding putative candidate genes (**Supplementary Table S1**, putative pseudogenes excluded), including the *Arabidopsis thaliana SUCROSE-PROTON-SYMPORTER 8 (AtSUC8)*, a candidate diversity-effect gene. The gene encodes for a proton symporter that is fueled by the electrochemical gradient across the membrane. *AtSUC8* is predominantly expressed in the root columella (Denyer et al., 2019; Graeff et al., 2021), and therefore in cells that are in direct contact with the soil, whose pH might affect its activity. To explore the idea that natural genetic variation at the *AtSUC8* locus could drive functional complementarity among *Arabidopsis* genotypes, we re-analyzed previously published data on competition between *Arabidopsis* genotypes (Wuest et al., 2019). Single individuals of ten tester genotypes (including Sav-0 and Uk-1) each competed separately with each genotype of a panel of 98 natural accessions, in a factorial design (**Figure 3 A and B**). For each tester-competitor pair, we determined specific combining abilities (SCAs) as in the present study (**Methods and Supplementary Figure 4 A and B**). We then tested for associations of these SCAs with between-genotype differences at single-nucleotide polymorphisms (SNPs) within the identified QTL on chromosome 2. After adjustment for multiple testing, only one SNP was significantly associated with a positive diversity effect within the QTL (**Figure 3C**, test for differences between mono-allelic and bi-allelic mixture SCAs by linear contrast t_947_ = 4.1; P = 5·10^−5^, Bonferroni-adjusted P = 0.007; standardized effect size = 3.2%). This SNP indeed resides in the *AtSUC8* coding region. Although this is not unequivocal proof that the identified SNP is the causal genetic polymorphism (it may instead be in tight linkage disequilibrium with the causal one), our finding provides further evidence that genetic differences in or around the AtSUC8 gene contribute to community overyielding in genotype mixtures.

**Figure 3:**
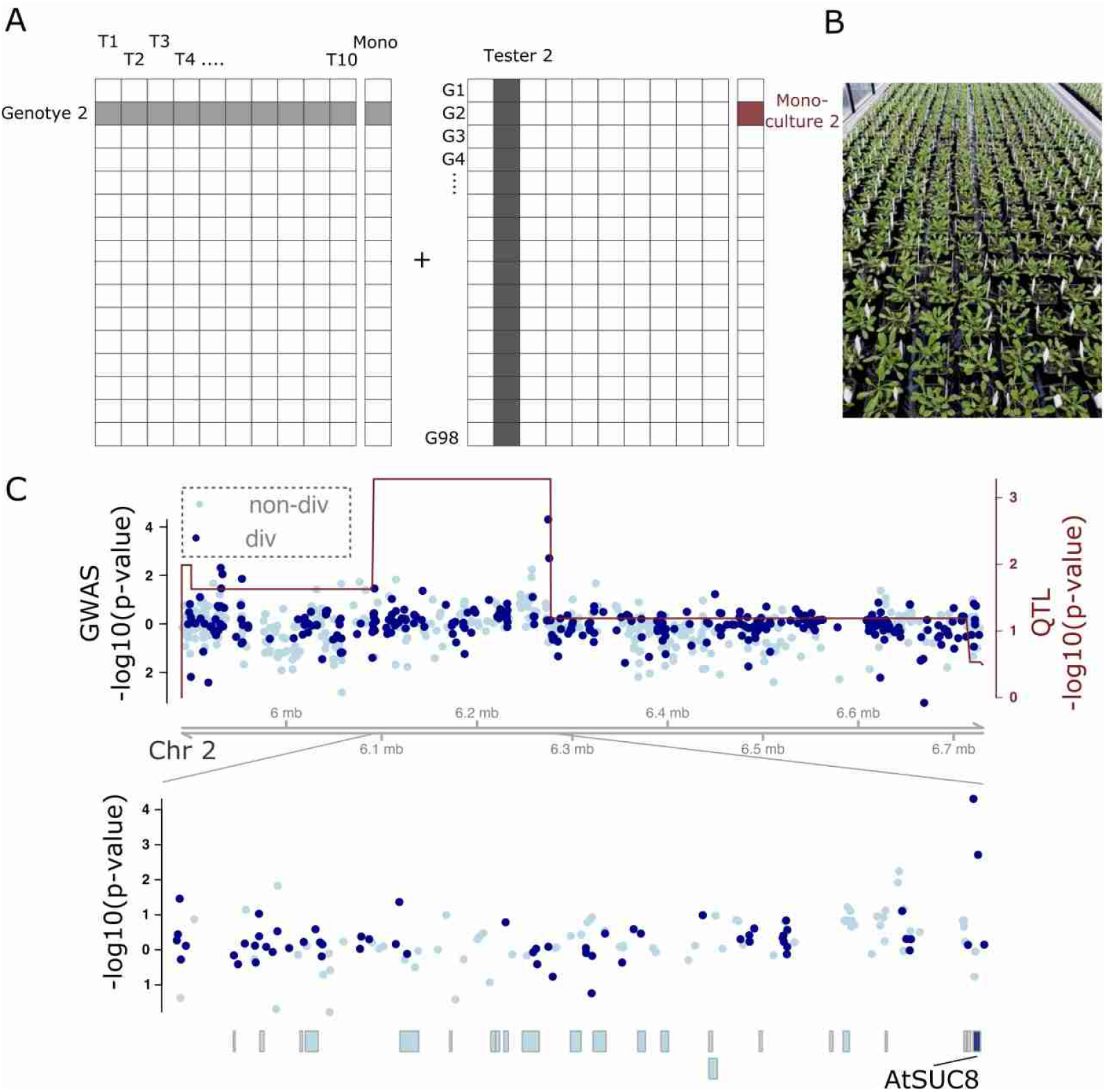
Single nucleotide polymorphism differences at the AtSUC8 locus associate with positive diversity effects in genotype mixtures. **A**. The experimental design represents a full-factorial combination of ten tester genotypes with each genotype of a panel of 98 natural Arabidopsis accessions **B**. Picture of the experiment **C**. The QTL mapping results (red line and right axis) overlaid with the genetic association results (blue dots and left axis). Light blue dots denote SNPs at which the Sav-0 and the Uk-1 tester lines do not differ (non-div), dark blue dots denote those at which they do differ (div). Dots above zero indicated positive diversity-SCA associations, dots below zero negative ones. Boxes in the bottom panel denote gene regions, the AtSUC8 gene region is colored dark blue.

SUC transporters are highly conserved within and across plant species. Sanger sequencing of the *AtSUC8* alleles from Uk-1, Sav-0 and the reference accession Col-0 confirmed the presence of several non-synonymous SNPs. Compared with the reference allele, the *AtSUC8* coding region of Sav-0 carries three amino acid replacements (one non-conservative), and the Uk-1 allele carries eleven amino acid polymorphisms (six non-conservative) (**Figure 4 A**). Among the latter, the K320T and the R472G replacements might be functionally relevant, because they also occur in the C24 accession which we had also used as tester genotype in the association study described above. C24 shares seven amino acid polymorphisms with Uk-1 and shows similar patterns of diversity effects across genotypes (**Supplementary Figure 4 C**). To determine whether the identified polymorphisms in Uk-1 and Sav-0 affect SUC8 function, we used sucrose uptake in a heterologous system as assay of function. We expressed the Uk-1 and Sav-0 variants of SUC8 in *Xenopus laevis* oocytes and measured their sucrose uptake kinetics. Whereas SUC8^Sav-0^ conferred efficient sucrose uptake as compared with mock-transformed oocytes, significantly lower import was observed with SUC8^Uk-1^ (**Figure 4 B**). We next tested if such functional protein differences also affect root growth under different pH conditions by growing 80 RILs from the Uk-1×Sav-0 RIL population on two media with pH ∼6.8 or ∼4.8. For this, we grew seedlings on these media and measured their root length. As expected, root length was reduced (by ≈50%) at low pH and (by ≈60%) in genotypes carrying BRX^Uk-1^ (**Figure 4 C**). Relative root length reduction at low pH versus neutral pH did not vary among genotypes carrying different *BRX* alleles (**Figure 4 C)**. However, the relative root length reduction was significantly smaller when genotypes carried the AtSUC8^Uk-1^ instead of the AtSUC8^Sav-0^ allele (linear model ANOVA F_1,74_ = 5.8; P = 0.02; **Figure 4 C**). These findings indicate that Uk-1 carries alleles at multiple loci, including *BRX* and *AtSUC8*, that change root growth and allocation in response to edaphic conditions, in particular environmental proton concentration. Overall, our results thus suggest that genetic differences associated with community overyielding in genotype mixtures are related to allele-specific differences in protein and root functioning.

**Figure 4:**
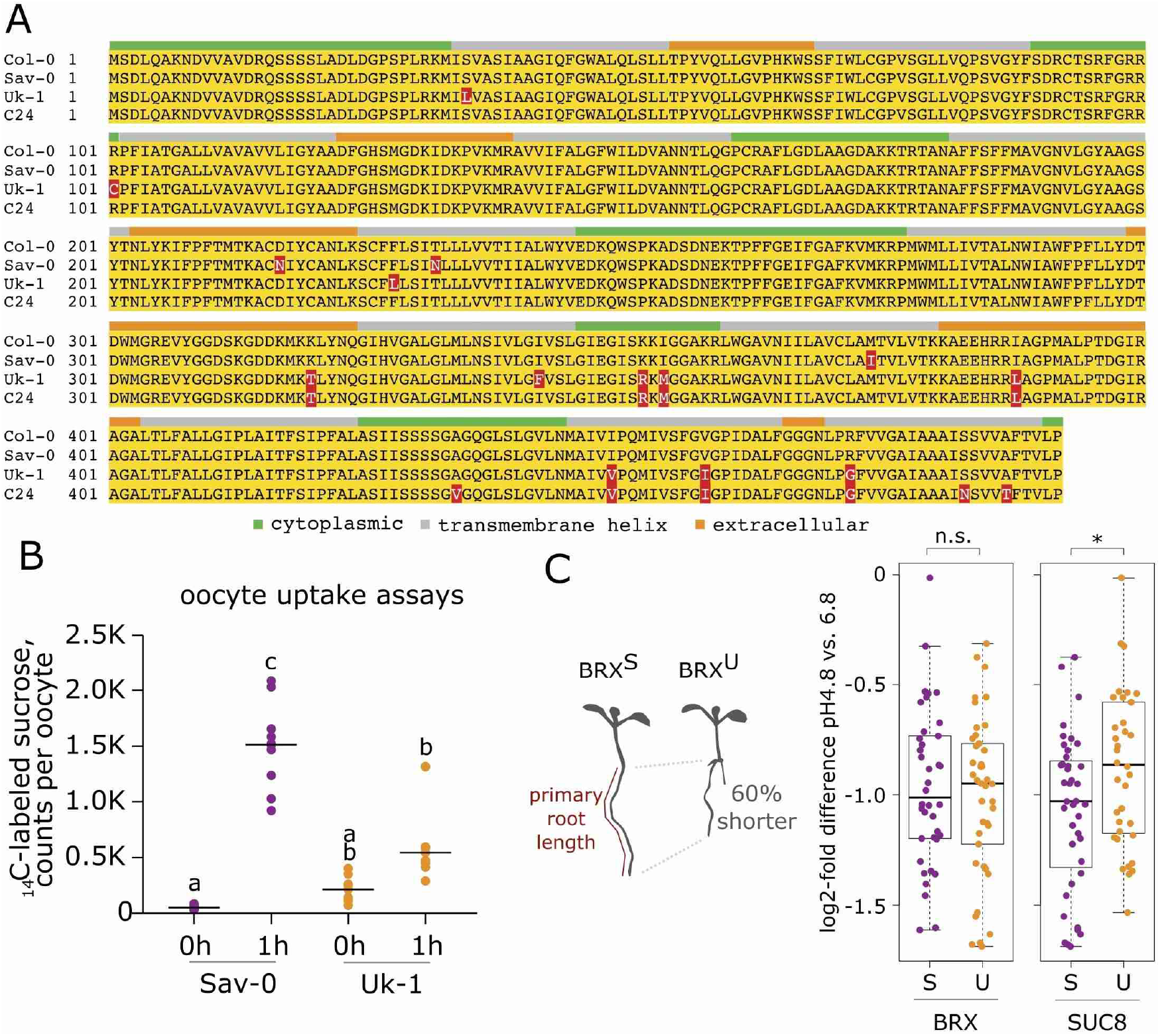
Genetic variation in AtSUC8 affects protein function and is associated with different root growth sensitivities to changes in substrate proton concentrations. **A**. Protein sequence alignments of natural SUC8 variants. Amino acid differences from Col-0 reference sequence are highlighted in red **B**. Sucrose transport activities of the Sav-0 and Uk-1 protein variants in oocytes. Different letters denote significant differences in Tukey’s post-hoc contrasts **C**. Primary root length differences of genotypes carrying carrying either Sav-0 (S) or Uk-1 (U) alleles at the two loci (BRX and AtSUC8), and grown on agarose plates exhibiting different substrate pH. Relative root length of different RILs carrying either alleles at the BRX (right) or AtSUC8 locus (left); shown are log2-fold root length differences of each RIL at pH 4.8 vs. 6.8 (e.g., a log-fold difference of –1 denoting roots being 2-fold shorter at pH 4.8 than at pH 6.8); * = p-value < 0.05; n.s. = not significant.

## Discussion

Here, we used two complementary genetic strategies, QTL- and association-mapping, to identify the genetic differences between *Arabidopsis* genotypes that overyield when grown in mixed-genotype communities. We found that a large proportion of the overyielding of mixtures of the *Arabidopsis* accessions Sav-0 and Uk-1 was due to allelic diversity at a major-effect QTL on chromosome 2. Two aspects of this QTL mapping study are worth noting. First, our QTL mapping resolution was very high despite using only 18 recombinant lines and their parents. This was due to the competition diallel experimental design in which genotypes with high-density marker maps are systematically combined into different communities. Second, although complex traits of individuals such as growth are often determined by genetic variants at many loci, each with small effect (Lynch and Walsh, 1998; MacKay et al., 2009; Wieters et al., 2021). Our results together with findings from recent studies (Wuest and Niklaus, 2018; McGale et al., 2020; Barbour et al., 2022; Montazeaud et al., 2022) suggest that complex community-level properties that depend on interactions between plant individuals can have surprisingly simple genetic underpinnings. Our work thus suggests that positive effects of plant diversity need not be irreducibly complex emergent properties but can have simple causes that are identifiable at the genetic level, even if the mixed genotypes differ at many positions along the genome. We think that understanding the origins of overyielding may in fact – at least in some cases – be simpler based on genetics than based on traits, where complementarity seems to generally manifest itself as a high-dimensional phenomenon involving a number of different traits (Montazeaud et al., 2020). The community genetic approaches presented here and elsewhere (Frachon et al., 2019; McGale et al., 2020; Turner et al., 2020; Sato et al., 2021; Subrahmaniam et al., 2021; Barbour et al., 2022; Montazeaud et al., 2022) may thus provide an effective way to understand the propagation of effects across different layers of biological organization, from genes to communities and ecosystems.

Identifying the genes that are important for ecosystem processes may ultimately also be useful to link ecological processes to some of the dominant evolutionary drivers (Johnson and Stinchcombe, 2007; Crutsinger, 2016). In our study, we were able to associate diversity at the *AtSUC8* locus with community-level overyielding. The respective gene encodes for a proton-sucrose symporter, i.e., a membrane-associated protein that utilizes a proton gradient to transport sucrose across membranes. The gene is expressed predominantly in root tissues that are in direct contact with the soil. Genetic differences at the *AtSUC8* locus affect protein function and were also associated with differences in root growth, in a substrate - pH dependent way. Soil chemistry, composition and texture and resulting effects on plant – plant interactions are major selective forces, but also important drivers of community structure (Tilman et al., 1997; McKane et al., 2002; Kahmen et al., 2006; Jiménez-Alfaro et al., 2018). Consistent with the idea that the Uk-1 genotype exhibits traits that make it better adapted to grow on acidic soil (Gujas et al., 2012), plants carrying the *AtSUC8*^Uk-1^ allele showed root growth that was less sensitive so substrate acidification. However, and perhaps surprisingly, genetic variation at the BRX locus itself, which had previously been shown to underlie adaptive divergence along this environmental gradient (Mouchel et al., 2004; Gujas et al., 2012), did not drive overyielding in our model communities. Future work should be able to establish possible reasons for these differences between *AtSUC8* and *BRX*, and the specific physiological and morphological effects of the identified genetic variation at the *AtSUC8* locus and their consequences for plant fitness under natural conditions.

One question that remains open is how specific aspects of SUC8-mediated trait differences account for overyielding in genetically diverse communities. We think that the different responses of root growth to changes in soil acidity associated with the *AtSUC8* locus promote the partitioning of the physical soil space between plants. In other words, these effects may result in different root foraging strategies in a substrate heterogeneous in soil solution pH, resulting in more efficient use of the available biotope space (Dimitrakopoulos and Schmid, 2004; Tylianakis et al., 2008; Jousset et al., 2011). A pH gradient, possibly at a very small scale, would then represent a niche dimension along which niche partitioning promotes community productivity. Obviously, there may be different environmental settings under which other traits, related to other genetic differences, may underly niche partitioning and complementarity among plants. In each case, the trait-based approaches currently applied for the study of ecological phenomena such as overyielding might strongly profit from gene-based approaches, ultimately not only at the within-but also at the between-species level. On the other hand, our work may offer new ways to design more sustainable cropping systems, in which species or genotype diversity can improve both yield and yield stability in the face of biotic and abiotic stress (Finckh et al., 2000; Zhu et al., 2000; Brooker et al., 2015; Litrico and Violle, 2015; Kristoffersen et al., 2020; Wuest et al., 2021). Here, the gene-centered approach may complement currently used trait-centered methods to facilitate the design of high-performing mixtures.

## Materials and Methods

### Germplasm

The Sav-0 and Uk-1 seeds were initially obtained from the Arabidopsis Biological Resource Center at Ohio State University. The Sav-0*Uk-1 RIL population was described previously (Mouchel et al., 2004). The lines used for the association analysis are described in detail in (Wuest et al., 2019)

### Plants and growth conditions

Seeds were sown directly on soil and germinated in trays covered with plastic lids under high humidity in a growth chamber at the University of Zurich Irchel Campus (16hrs light, 8 hrs dark; 20°C, 60% humidity). The soil substrates are described below. After approximately two weeks, the trays were moved into a greenhouse chamber, where day-time and night-time temperatures were maintained around 20–25 °C and 16–20 °C, respectively. Additional light was provided if required to achieve a photoperiod of 14–16 hours. Seedlings were thinned continuously until a single healthy seedling remained per position. The pots were watered *ad libitum*, and in case of high herbivory pressure by larvae of the dark-winged fungus gnat the insecticide ActaraG (Syngenta Agro AG) was applied according to the manufacturer’s recommendation. The date of harvesting was determined through the occurrence of 5– 10 dehiscent siliques on the earliest flowering genotypes in a given block. The aboveground biomass was dried at 65°C for at least three days and then weighed.

### Assessing accession pair mixtures

Nine accession pairs, for which recombinant inbred line populations are publicly available, were chosen for the screen of pair-wise interactions through comparisons of monoculture and two-genotype mixtures. A further pair was chosen based on a large estimate of mixture effects in a previous study. These selected genotypes were grown as either monocultures or pair-wise mixtures on different soils and in pots of different size as follows: peat-rich Einheitserde ED73 soil substrate (pH ∼5.8, N 250 mg L^-1^; P_2_O_5_ 300 mg L^-1^; 75% organic matter content; Gebrüder Patzer GmbH, Sinntal-Jossa, Germany) and in 6*6*5.5 cm or 7*7*8 cm or 9*9*10 cm pot sizes, a 4:1 mixture of quartz sand:ED73 and 7*7*8 cm pots, and *Arabidopsis* legacy soil, i.e., soil collected from an unrelated previous experiment on which *Arabidopsis* had grown (originally ED73). Each monoculture or mixture composition in each soil or pot size was grown in each of seven blocks, with the exception of communities on sand-rich and legacy-soil conditions. The legacy and sandy soil conditions were included only in five of the blocks for logistical reasons. Community overyielding in genotypic mixtures containing Sav-0 and Uk-1 was confirmed by growing either i) four plants in medium sized pots (7*7*8 cm); ii) four plants in small pots (5.5*5.5*6 cm) or iii) two plants in small pots, all containing ED73 soil. For each pot/density type, 48 mixtures and 24 of each monoculture were sown, treated and processed as described above.

### QTL mapping and association study

The QTL-mapping experiment was designed as a half-diallel containing all pair-wise combinations, and monocultures of, 18 RILs derived from Sav-0 and Uk-1 (Mouchel et al., 2004) and the two parents. The experiment was performed in four sequential blocks; we used a soil consisting of 3 parts ED73 and 1 part quartz sand for the first two blocks. However, because seedling establishment was rather poor on this soil, we changed soil type in blocks three and four to 1 part ED73 and 3 parts sand. Plants were grown and harvested as described above (42–51 days after sowing).

Experimental conditions for the genome-wide association experiment are described in detail elsewhere (Wuest et al., 2019). In short, the association study experimental design consisted of a full factorial competition treatment of growing ten tester genotypes (Sav-0; Uk-1; Col-0; Sf-2; St-0; C24; Sha; Bay-0; Ler-1; Cvi-0) with each genotype of an association panel of 98 natural *Arabidopsis* accessions (a subset of the RegMap population (Horton et al., 2012), including all monocultures and in two replicate blocks. Each community consisted of two plants (one plant per genotype). The raw data of the association study are available at https://zenodo.org/record/2659735#.YCt0u2Mo8mI).

### Genotyping and line re-sequencing

For the 18 RIL genotypes used in the QTL-mapping competition diallel, we performed whole-genome resequencing and genotype reconstructions before the genetic analysis. DNA extractions for genome resequencing, library preparation, sequencing and genome reconstruction was performed as previously described (Wuest and Niklaus, 2018), whereby the genome reconstruction approach broadly followed the method described by Xie and colleagues (Xie et al., 2010). Raw reads of resequencing the parental accessions Sav-0 and Uk-1 were downloaded from the NCBI SRA homepage (www.ncbi.nlm.nih.gov/sra, SRX011868 and SRX145024). To genotype a wider set of RIL lines at the *AtSUC8* locus (At2g14670), a Cleaved Amplified Polymorphism (CAPS)-marker assay was developed based on a EcoRV-restriction site in the SUC8 coding sequence that is present in the Sav-allele but missing in the Uk-Allele using PCR primers 5’-GGA GAG TGT TGT TAG CCA CGT C-3’and 5’-ACG ATG TGG TAG CTG TAG ATA GAC-3’. DNA extractions for CAPS-genotyping were performed using the protocol following Edwards and colleagues (Edwards et al., 1991). For four RIL genotypes where the PCR-genotyping yielded ambiguous results, so we inferred it from flanking markers AtMSQTsnp 123: (Chr 2 pos 1798324) and AtMSQTsnp 138 (Chr 2 pos 8370574) (Kim et al., 2007). We also tried to identify RIL-lines that exhibited heterozygosity at the *AtSUC8* locus to isolate heterogeneous inbred families, but failed to find any among the 101 lines screened.

To verify polymorphisms identified in the resequencing, Sanger sequencing of the *AtSUC8* alleles was performed by amplifying the gene body from genomic DNA using oligonucleotides 5’-ATG AGT GAC CTC CAA GCA AAA AAC GAT-3 and 5’-TTA AGG TAA CAC GGT AAA TGC CAC AAC ACT GC-3’. The PCR fragments were then sequenced using those same oligonucleotides as well as oligonucleotide 5’-CAC AAT GAC TAA AGC ATG TGA C-3’. The C24 allele of SUC8 was retrieved from published sequence data (Jiao and Schneeberger, 2020). Note that because of genomic rearrangements, the gene ID for *AtSUC8* (AtC24-2G29550) in the C24 accession differs from the other accessions.

### Oocyte uptake assays

Oocyte assays were performed essentially as described (Fastner et al., 2017). Briefly, the SUC8 cDNAs were cloned into pOO2 (Ludewig et al., 2002). cRNA was synthesized using the mmessage mmachine kit (Lifetechnologies). Oocyte s were injected with 50 nL of 150 ng/μL cRNA and incubated in Barth’s (88 mM NaCl, 1 mM KCl, 2.4 mM NaHCO_3_, 10 mM HEPES-NaOH, 0.33 mM Ca(NO_3_)_2_ x 4 H_2_O, 0.41 mM CaCl_2_ x 2 H_2_O, 0.82 mM MgSO_4_ x 7 H_2_O pH7.4) for four days. For uptake experiments 10 oocytes were kept in 1 ml Barth solution supplemented with[^3^H]-sucrose or [^14^C]-sucrose at a final concentration of 1 mM or substrate-free control for one hour. Afterwards, Oocytes were washed twice in Barth solution containing Gentamycin and were then separated into scintillation vials. 100 μl of 10 % SDS (w/v) was added to each scintillation vial and the samples were incubated for 10 minutes. Then 2 mL of scintillation cocktail (Rotiszint eco plus, Roth, Germany) was added and the vials were vortexed vigorously. Radioactivity was determined by liquid scintillation counting. Experiments were carried out using [^14^C]-sucrose and repeated with [^3^H]-sucrose yielding essentially identical results. [^14^C]-sucrose (536 mCi/mmol, 1 mCi/ml) and [^3^H]-sucrose (3 Ci/mmol, 1 mCi/ml) were purchased from Hartmann Analytic, Braunschweig, Germany)

### Plate assays and root measurements

Seeds were surface-sterilized with 70% ethanol, followed by 15 minutes in a solution containing 1% bleach and 0.01% Triton-X100 and three sequential washes, then left for stratification at 4°C overnight. Square MS plates (12 cm) were prepared with 0.8% agarose (instead of agar) and containing 1% sucrose (w/v). The pH was adjusted to 4.5 or 7 using hydrochloric acid or potassium hydroxide and the medium autoclaved. After autoclaving, the measured media pH was again determined (4.8 and 6.8). Six seeds of each of six different genotypes were sown on a plate pair (identical sowing pattern on pH 4.8 and 6.8) and grown in a climate chamber with long-day conditions (16 hours light at 20°C; 8 hours dark at 16°C) for seven days. Plates were scanned twice, once after 3 days and again after 7 days using an EPSON flatbed scanner (model 2450). The primary root length of seedlings was measured using the Fiji software (Schindelin et al., 2012).

### Statistical analyses

In the screen for consistently positive pairwise interactions between genotypes, we fitted a linear model of community biomass as function of genotype composition and substrate type (i.e., substrate composition or volume), including a block term. Overyielding of a genotype pair on a given substrate was then estimated as linear contrast between the average monoculture productivity and the mixture productivity (i.e., specifying the contrast matrix K=[-0.5, −0.5, 1], equivalent to the term 1m_AB_ – 0.5m_AA_ – 0.5m_BB_ for the case of a monocultures and mixtures of genotypes A and B), using the glht-function of the multcomp-package (Hothorn et al., 2008).

The mapping experiment was performed on two different substrates (two replicated blocks each), and both mean and variance of community productivities differed across substrates. The blocks with more nutrient-rich substrate also had some pots with missing plants due to seedling mortality, which were removed for the analysis. In order to combine all four blocks for the estimation of specific combining abilities, we therefore first estimated mean community biomass within substrate and calculated specific combining abilities (SCA) within substrates from average total pot biomass values (BM) as BM = Z*u + SCA whereby Z is the design matrix describing genotype composition of a mixture. To make SCAs comparable across substrates, we divided SCA through the mean pot biomass produced on this substrate. The standardized SCA_ij_ value of a genotype composition (containing genotypes G_i_ and G_j_) was then estimated by averaging across substrates. SCA outliers were removed if they differed more than two standard deviations from the population mean in their absolute value. QTL mapping of standardized mixture SCA estimates was then performed by a marker regression approach, where we first fitted a linear model predicting *SCA from allelic composition (3 levels, SS, UU, SU)*, followed by a contrast between allelic monocultures and mixtures (e.g., SCA_SU_ – 0.5(SCA_UU_ + SCA_SS_), again using the glht function

A LOD score (-log10(p-value) of 3 was considered significant, as determined by large-scale simulations (Van Ooijen, 1999) assumptions: two QTL genotypes, “bi-allelic” and “mono-allelic” and an average chromosome length of 200 cM for *Arabidopsis* genotype pairs, where recombination events are combined in communities). Such a threshold is also in agreement with our previous work comparing this approach to a standard QTL mapping method and a LOD-cutoff based on re-sampling (Wuest and Niklaus, 2018).

### Analysis of association-study competition experiment

The association study represents a factorial design in which each of ten different genotype (testers) was grown in combination with each of 98 different *Arabidopsis* genotypes, with all monocultures realized too. This design was replicated in two blocks. Pots with missing data (e.g., due to seedling mortality) were removed from the analysis. A genotype’s general combining ability was estimated as described above within each block and values were then averaged across blocks.

Pot biomass depended non-linearly on average genotype GCA (Supplementary Figure 4). To determine SCAs, we therefore used a quadratic form of the mean GCA to adjust for this non-linearity. Marker regressions on these SCA values for the SNPs within the QTL interval were performed as described for the QTL mapping approach described above.

## Author contributions

SEW conducted and analyzed the screen for overyielding amongst *Arabidopsis* accessions with support from ME, and the QTL mapping experiment, with support from BS and PAN. CSH provided the genotyped Uk-1 x Sav-0 RIL population. SEW and NP performed the association study with input from UG. SR and CSH performed the sequence analyses of the *AtSUC8* gene. LS and UH conducted the oocyte uptake assays. SEW and UG raised funding. SEW together with PAN wrote the first draft of the manuscript. All authors revised and approved the final version of the manuscript.

## Acknowledgements

We thank Matthias Philipp, Daniela Stöckli and Nicole Ponta for help with plant maintenance and measurements and Matthias Furler for technical support in the greenhouse. This work was supported by the University of Zurich, Agroscope, grants from the Swiss National Science Foundation (Ambizione Fellowship PZ00P3_148223) and the University Research Priority Program “Evolution in Action” and of the Swiss National Science Foundation to S.E.W., and Advanced Grant of the European Research Council (AdG #250358) to U.G..

## Data availability

The datasets described are available through the Zenodo data repository (DOI:10.5281/zenodo.7104830).

## Supplementary Figures

**Supplementary Figure 1.**
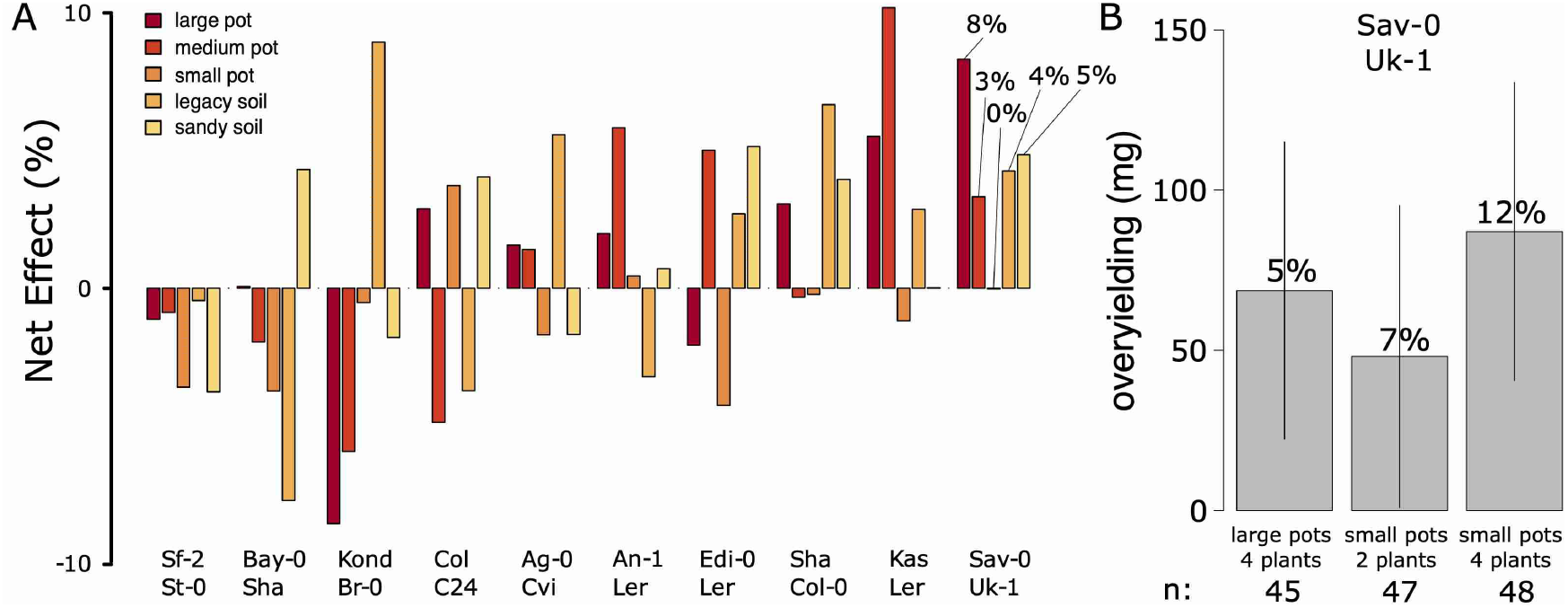
**A**. A screen for consistent genotypic diversity effects between divergent *Arabidopsis* accession pairs. Shown are estimates of net overyielding (observed mixture yield compared with average yields of component monocultures) of ten *Arabidopsis* accession pairs across different soil types or pot sizes. For each estimate, seven pots (large pot, medium pot, small pot) or five pots (legacy soil, sandy soil) of each monoculture and the mixture were sown, resulting in a total of 930 pots containing four plants each. Note that both consistent negative (left) or consistent positive (right) effects appear. Furthermore, a soil-by-diversity interaction in the Bay-0 * Sha combination has been examined in more detail previously (Wuest and Niklaus, 2018). **B**. Confirmation of consistently positive genotypic diversity effects in the genotype combination Slavice-0 (Sav-0) and Umkirch-1 (Uk-1) under three different conditions. Shown are estimated net overyielding for each condition, number above bars indicate the relative net effect (%). Error bars: +/-s.e.m.

**Supplementary Figure 2.**
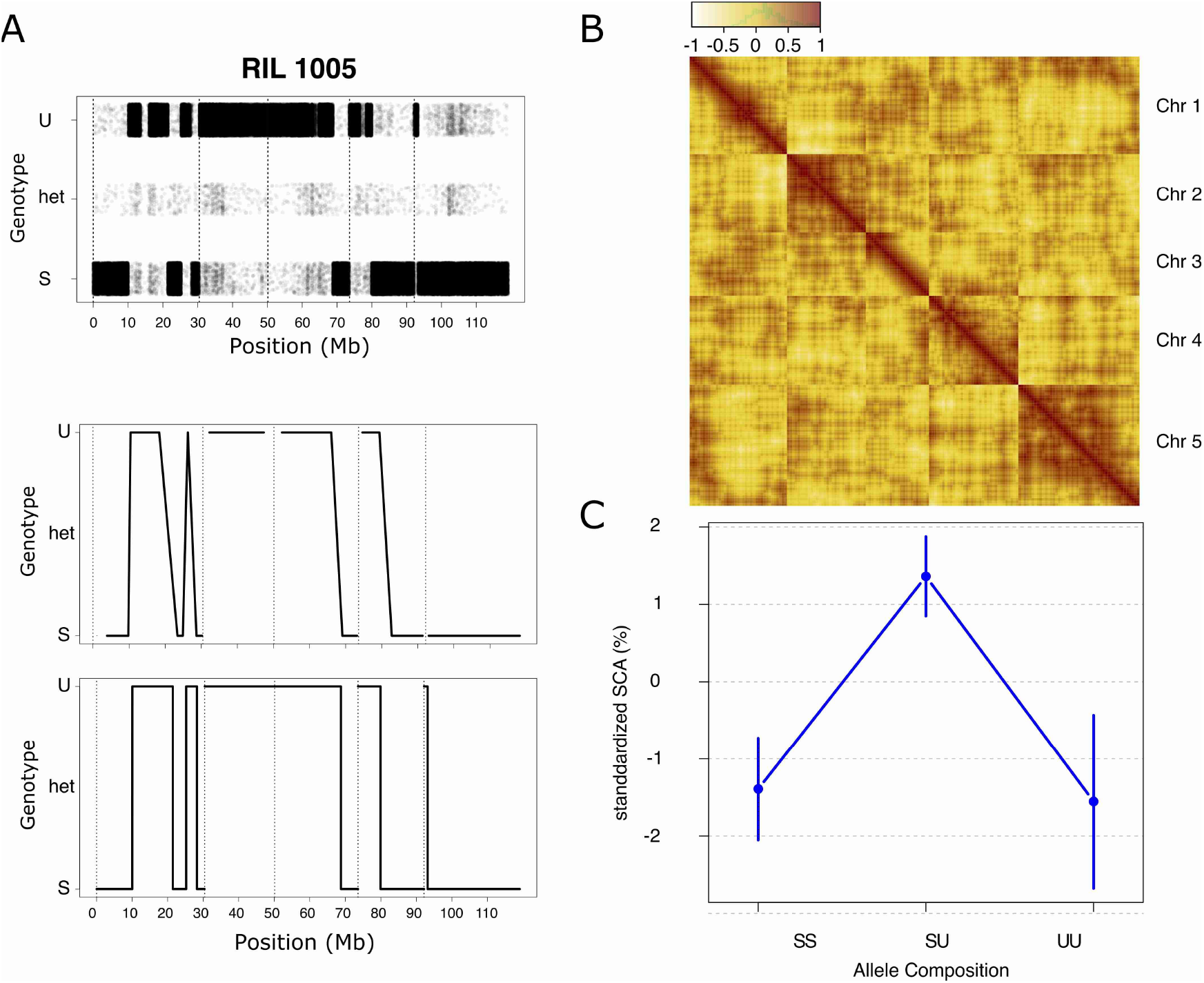
Reconstruction of RIL genotypes from low-coverage genome re-sequencing and QTL effect sizes. **A**. Top: Genotype calls across the genome in RIL US1005; and comparison of molecular markers (middle) and genotype reconstruction based on low-coverage genome re-sequencing (Viterbi-Path, bottom). **B**. Correlations of allelic compositions between all markers and across all genotype combinations **C**. Effect of allelic composition on specific combining abilities at the QTL chromosome 2 (QTL2, bottom).

**Supplementary Figure 3.**
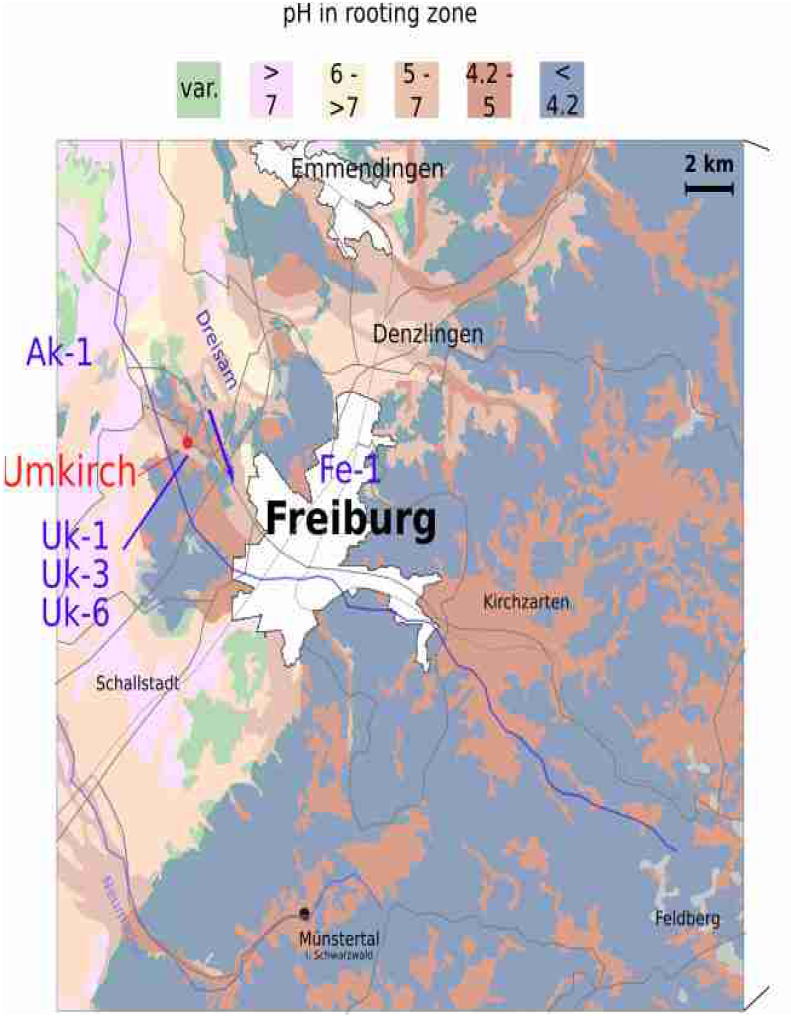
Soil acidity map of the southern black forest region, the area in which the Uk-1 accession was collected. Transect sampling performed by Shindo and colleagues (Shindo et al,): purple arrow. Data from http://maps.lgrb-bw.de/. var = variable

**Supplementary Figure 4:**
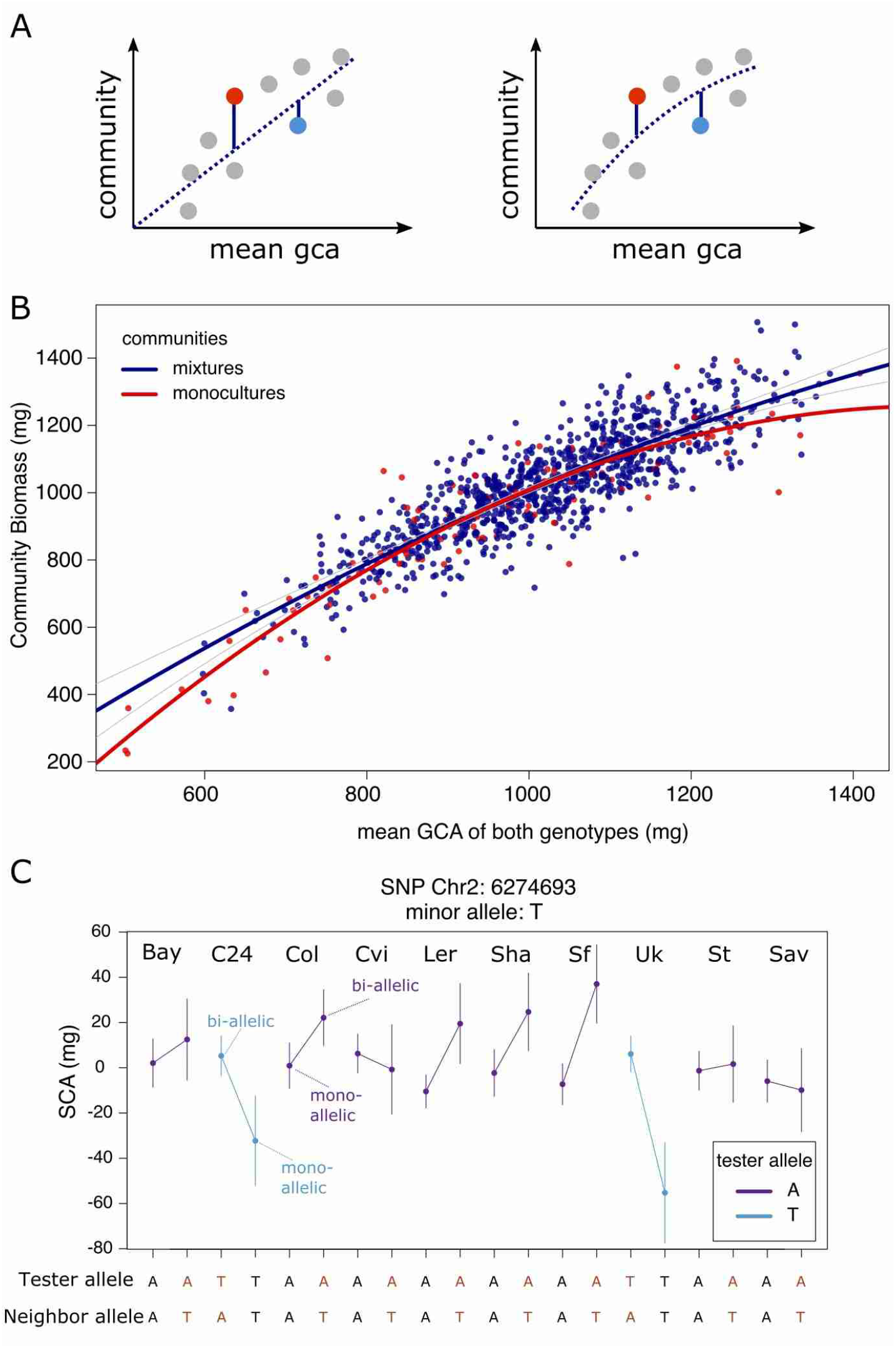
Determination of SCAs in factorial (tester-associate) competition design for GWAS and SCA across different tester lines and the different allelic diversity levels at a SNP within *AtSUC8*. **A**. Specific combining ability of a genotypic composition is typically estimated from deviates of observed community productivities from expectations (in this case, the average GCA of both genotypes); however, because different communities varied so strongly in total productivities, the relationship between the mean GCA of a genotype composition and the overall community productivity might become non-linear (e.g., driven by increasingly restricted space for combinations of highly productive genotypes). In this case, such a systematic relationship can first be modeled, and the SCA estimated as the deviation from this modeled relationship. **B**. Observed relationship between the average GCA of a genotype composition and its community productivity. **C**. Uk-1 and C24 both carry the minor (T) allele at SNP Chr2-6274693. When combined with genotypes also carrying the minor allele, the resulting mixtures show on average lower SCA, when combined with genotypes carrying the major allele (A), they exhibit on average higher SCA.

**Supplementary Table 1:** Descriptions of protein-coding genes found within the QTL on chromosome 2.

| Locus | Description | Symbols |
| --- | --- | --- |
| AT2G14378 | Encodes a ECA1 gametogenesis related family protein | NA |
| AT2G14390 | Hypothetical protein | NA |
| AT2G14440 | Leucine-rich repeat protein kinase family protein | NA |
| AT2G14460 | hypothetical proteinH | NA |
| AT2G14500 | F-box family protein | ATFDB14 |
| AT2G14510 | Leucine-rich repeat protein kinase family protein | NA |
| AT2G14520 | CBS domain protein (DUF21) | NA |
| AT2G14530 | Encodes a member of the TRICHOME BIREFRINGENCE-LIKE gene family | TBL13 |
| AT2G14540 | Serpin 2 | SRP2; ATSRP2 |
| AT2G14560 | Encodes LURP1, a member of the LURP cluster (late upregulated in response to *Hyaloperonospora parasitica*). LURP1 is required for full basal defense to *H. parasitica*. | NA |
| AT2G14580 | Pathogenesis related protein, encodes a basic PR1-like protein. | PRB1; ATCAPE7; ATPRB1 |
| AT2G14610 | PR1 gene expression is induced in response to a variety of pathogens. It is a useful molecular marker for the SAR response. Expression of this gene is salicylic-acid responsive. | PR1; ATCAPE9 |
| AT2G14620 | Xyloglucan endotransglucosylase/hydrolase 10 | XTH10 |
| AT2G14635 | ARABIDILLO protein | NA |
| AT2G14660 | Thymocyte nuclear-like protein | NA |
| AT2G14670 | Sucrose-proton symporter 8 | SUC8; AtSUC8 |

